# Microbial source tracking of human and animal fecal contamination in Ecuadorian households

**DOI:** 10.1101/2025.08.22.671888

**Authors:** Kelsey J Jesser, Viviana Alban, Aldo E. Lobos, Javier Gallard-Góngora, Gabriel Trueba, Gwenyth O Lee, Joseph NS Eisenberg, Valerie J Harwood, Karen Levy

## Abstract

Exposures to both human and animal feces pose human health risks, particularly for young children in low- and middle-income country (LMIC) settings where domestic animals are common, water and sanitation infrastructure is often limited, and enteropathogen transmission is high. Microbial source tracking (MST) markers specific to feces from humans and particular animal types can be used to identify the provenance of microbial contamination, yet most MST studies explore few household environmental sample types, limiting understanding of how marker utility varies by matrix. We validated qPCR assays for six MST markers and quantified their prevalence in 585 samples from 59 households spanning an urban–rural gradient in northwestern Ecuador. We used GenBac3 to test for general fecal contamination, and HF183, Rum2Bac, Pig2Bac, DG37, and GFD to test for human, ruminant, swine, dog, and avian contamination, respectively. Approximately 10 sample types were collected per household, including: rinses of child and adult hands, swabs of floors and surfaces, soil, domestic and drinking water, and food. GenBac3 and HF183 were detected in 77.82% and 15.36% of samples, respectively. Animal-associated markers were detected less frequently, in 0.5–4.1% of samples. However, when present, animal marker concentrations were comparable to HF183. Host-associated markers were most often detected in adult and child hand rinse and floor samples, and GenBac3 concentrations were highest in hand rinses. HF183 detection on adult caregiver hands was associated with increased odds of HF183 detection on children’s hands and floors.

**Importance:** Understanding the sources and pathways of detectable household environmental fecal contamination is critical for identifying how exposures occur and for developing targeted interventions to reduce risk of enteric infection By linking contamination on caregiver hands to that on children’s hands and floors, we highlight a likely route for pathogen transfer in the home. The inclusion of multiple host-associated markers across a wide range of sample types reveals patterns that narrower studies may miss, offering new insights into the complex ecology of fecal contamination. These findings can inform sampling strategies, guide risk assessments, and support the design of interventions aimed at reducing child exposure to enteric pathogens in similar high-risk settings.

## INTRODUCTION

Contamination of the environment with feces due to inadequate access to water, sanitation, and hygiene (WaSH) infrastructure contributes to a high burden of diarrheal disease in low- and middle-income countries (LMICs)^1,2^. In addition to acute diarrhea, persistent and recurring enteric infections, particularly in children under age five, are associated with serious negative health outcomes, including growth shortfalls and impaired cognitive, gut microbiome, and immune development^3–5^. Efforts to reduce enteric disease burdens have primarily focused on reducing exposures to human fecal contamination through increased latrine coverage, hand washing, and other interventions. However, recent trials have demonstrated that these interventions are insufficient to reduce human exposures to enteric pathogens or improve health^e.g., 6–10^. Other exposure pathways, and in particular the role of animals as contributors to reservoirs of fecal contamination and the transmission of enteric pathogens, have been increasingly recognized^11–15^. Animal feces harbor high concentrations of potentially zoonotic pathogens, such as *Salmonella* and *Campylobacter*^16–20^, and human exposures in the domestic environment may occur through direct contact, food contamination, and/or contamination of water sources, soil, and surfaces^12,21–24^. It is important to quantify reservoirs of environmental fecal contamination and differentiate between human and animal sources to understand and effectively interrupt enteric disease transmission.

Direct detection of enteric pathogens in the environment is difficult due to the diversity and low concentrations of pathogens in environmental samples^25,26^. For this reason, assessments of environmental exposures to enteric pathogens commonly focus on molecular or culture-based detection of fecal indicator bacteria (FIB), including *Escherichia coli*, fecal coliforms, and enterococci^23,27^, which are abundant members of human and animal fecal microbiomes and are found at high concentrations in sewage and feces^27^. Many studies have reported high FIB concentrations in environmental samples from LMIC household settings, including water, soil, hands, and surfaces^12,28–31^. However, FIB are imperfect indicators of environmental fecal contamination as they can originate from soils and other non-fecal sources, do not correlate well with pathogen concentrations, and do not differentiate human versus animal contamination^27^.

In contrast, molecular microbial source tracking (MST) markers can quantitatively measure and discriminate between human and animal sources of environmental fecal contamination. Quantitative PCR (qPCR) assays for MST markers target fragments of genes from host-associated enteric bacteria or viruses in environmental samples. Most MST assays were designed for use in high-income countries, where the gut microbiota is distinct from populations in LMIC settings due to differences in diet and lifestyle^32^. Because variability in human and animal gut flora can impact marker performance (both sensitivity and specificity), it is essential to validate MST markers for use in a given geographical and cultural setting.

Several studies have validated and applied MST methods to measure and differentiate between sources of fecal contamination in household and/or community water sources, hand rinses, soil, and other environmental sample types in LMICs, including Bangladesh^21,33,34^, India^35,36^, Nepal^37^, Mozambique^23^, and Peru^24^. The results of these studies confirm the ubiquity of human and animal fecal contamination in LMIC environments and have highlighted the likely contributions of animal feces in limiting the success of intervention trials. However, these previous efforts have focused on relatively few sample types and MST markers. In 12 studies published between 2015 and 2025, there was an average of 2.33 household sample types and 3.00 markers used per study^21,23,24,33–41^. The most commonly analyzed sample types were hand rinses (10 studies), drinking water (9 studies), soil (9 studies), and floors (6 studies), with foods, objects, and surfaces samples examined less often. In addition, several studies have reported poor marker performance (frequently defined as <80% sensitivity and/or specificity) for MST validation efforts in LMIC settings ^e.g., 23,37,42–44^.

Here, we report the results of MST marker validation and application to measure the sources and quantities of household fecal contamination in the LMIC setting of northwestern Ecuador. We conducted this study in communities located along an urban-rural gradient. As we have previously reported, these communities encompass a range of lifestyles, WaSH conditions, and interactions with animals^45,46^. We targeted animal-owning households and collected up to 10 types of samples per household, including water, surfaces, hands, food, and soil. All sample types were tested with five validated general, human, and animal-associated MST markers. Our objectives were twofold: i) to identify which household sample types most frequently contained detectable fecal contamination, and ii) to determine the most prevalent and abundant sources of fecal contamination within households.

## METHODS

### Study design

This study was conducted in communities that span an urban-rural gradient in the high enteric pathogen transmission setting of northern Ecuador. The inclusion of multiple sites with varying urbanicity offered variability in animal interactions and exposures, as we have reported previously^47^. Sites included the ‘urban’ city of Esmeraldas (population ∼150,000), the ‘intermediate’ town of Borbón (population ∼5,000), and several ‘rural’ communities near Borbón (populations ∼400-920). We enrolled *n=*58 households that had previously reported animal ownership (Table S1). We focused on households with that owned animals to increase our chances of detecting animal-associated markers and thus enhance our understanding of what sample types we should use in future studies to detect animal sources of fecal contamination in both animal-owning and non-owning households. Approximately half of enrolled households (*n=*31) had a child aged 6-24 months in the home. Figure 1 provides an overview of the study design.

**Figure 1:**
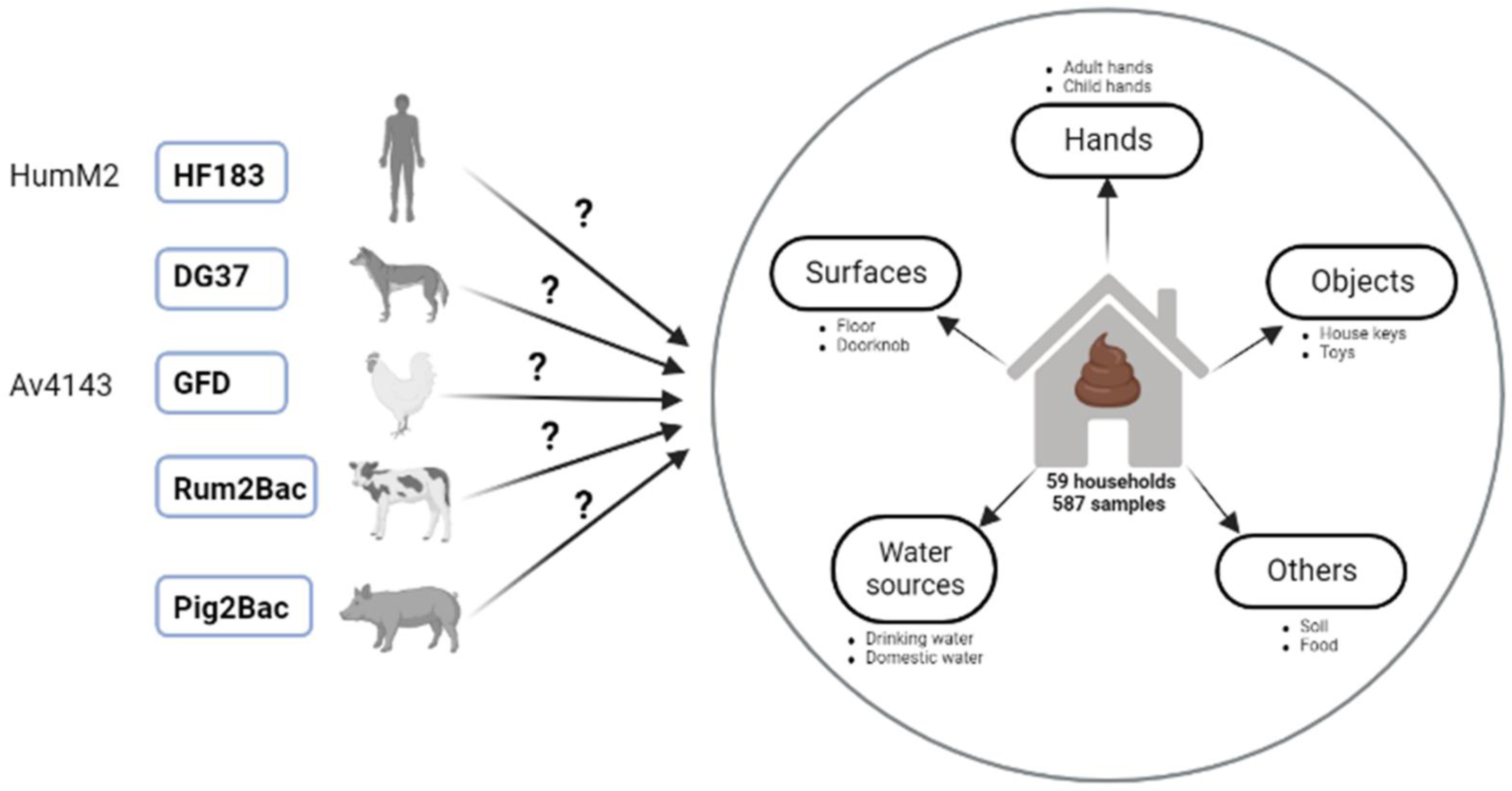
Summary of study design. Candidate qPCR assays tested for sensitivity and specificity in the study region are shown on the left. Assays selected for use in the study following MST marker validation are shown in bold typeface with blue boxes. These selected markers were used to test DNA isolated from a variety of household samples for human and animal fecal contamination.

### Ethics

The study was approved by the institutional review boards of Emory University (IRB00101202), the University of Washington (IRB0010343 and 00014270), and the Universidad San Francisco de Quito (2018-022M and 2021-089M). The protocol was also reviewed and approved by the Ecuadorian Ministry of Health (MSPCURI000253-4).

### Microbiological analyses

#### MST validation study

Prior to application for household samples, we validated MST markers for specificity (reactivity only with feces from target host), sensitivity (frequency of detection in target hosts), and concentration in target host feces. Animal feces (n=80 from five animal types: dogs, birds/poultry, pigs, and ruminants), human feces (n=7), and human sewage samples (n=5) were collected from in and around households in the study region and frozen immediately in a liquid nitrogen dewar. Samples were transported to the Universidad de San Francisco de Quito (USFQ) laboratory facilities, where DNA was extracted from 250 mg feces or 250 mL sewage using the ZymoBIOMICS DNA Miniprep kit (Zymo Research, Irvine, CA) according to the manufacturer’s protocol. DNA was eluted in 100 μL of Zymo DNA Elution buffer and stored at −80°C. Extracts were hand-carried from Quito to the University of South Florida (USF). Gel ice packs (TECHNI ICE, Frankston, Victoria, Australia) to maintain cold chain during the approximately 12h travel time. Extracts were verified to have remained frozen upon receipt at USF and stored at −80°C prior to analysis. DNA concentration was measured both before freezing at USFQ and prior to qPCR analyses at USF using a Qubit fluorometer to ensure DNA had not degraded during transport (Thermo Fisher).

#### Household environmental sampling

We aimed to collect approximately *n=*10 samples per household, resulting in *n=*585 total samples. Sample types encompassed aliquots of drinking water and domestic water, rinses of child and adult hands, swabs and rinses of surfaces (floors, dish drying and food preparation areas, doorknobs) and objects (TV remotes, keys, cell phones), food, and soil. Sample numbers are available in Table S2. Household data, including information on animal ownership and household sample characteristics (e.g. floor type, water source, hand cleanliness, etc.) were obtained in ODK during the same visit using surveys and observations and are summarized in Table S3.

##### Water samples

Depending on the volume the household was able to provide, either 500 mL or 1 L of drinking and domestic water samples were collected in 1 L Whirl-pak bags (NASCO WHIRIL-PAC®, USA). Before collection, we determined the presence of chlorine residues using Mquant^TM^ Chlorine Test Strips (MilliporeSigma). If chlorine was detected, we used Whirl-pak bags with sodium thiosulfate for sample collection to neutralize the chlorine.

##### Hand and object rinse samples

Child and adult hand rinses were collected by asking an adult caregiver to insert their hands or their child’s hands into Whirl-pak bags containing 200 mL sterile 1M phosphate buffered saline (PBS). Participants were asked to remove any jewelry prior to sampling. Hands were massaged while submerged in PBS for 60 seconds, with effort made to rub each finger, scrape under the fingernails, and massage the entire palm and back of the hand. Rinses of objects (toys and keys) were collected in the same way if the objects were available.

##### Swab samples

Swabs of floors, dish drying surfaces, and food preparation surfaces were taken using a 30 cm x 30 cm stainless-steel surface template disinfected with 10% bleach followed by 70% EtOH. Sampling locations were based on responses to survey questions about where the family typically spends time, cooks, or dries dishes. The template was placed on the surface, and a sterile cotton swab was wetted in a microcentrifuge tube containing 1 mL of sterile PBS. We swabbed within the template in three directions (vertically, horizontally, diagonally) to ensure complete coverage. The swab was clipped with sterilized scissors into a 2 mL microcentrifuge containing 1 mL Zymo DNA/RNA Shield. The surface was then swabbed a second time in three directions with a dry swab to collect any remaining wetting solution. The second swab was clipped into the same tube.

Doorknob surface and object swab sampling for TV remotes and cellphones was conducted in the same way, except that we aimed to swab the entire object. Doorknob sampling was based on the door household residents indicated was most frequently used. Keys and cell phones were sampled if they were available.

##### Food samples

We called households the day before environmental sampling and asked residents keep food (cooked rice, green plantains, or colada) prepared and stored using their usual practices. Household residents were asked to place stored food into a sterile Whirl Pak bag using their typical serving method (e.g. using a utensil or fingers).

##### Soil samples

Soil from the household compound was collected by marking the area to be sampled with a 10% bleach and 70% EtOH-sterilized 30 cm x 30 cm surface template. A metal scoop that was also surface-sterilized with 10% bleach and 70% EtOh was used to gently disrupt the top 1-2 cm of soil within the template area, and approximately 50 g of unsieved soil was collected in a Whirl-Pak bag. Soil sampling location in the yard or compound was selected based on responses to survey questions about where the household, and especially young children, spend time.

##### Field blanks

Two field blanks were collected each day that samples were collected. The first was a sterile Whirl-Pak bag filled with 200 mL PBS that was opened in a household for approximately the time needed to collect a hand or object rinse sample. The second was a sterile swab that was opened in a household, held in the air for approximately the amount of time needed to collect a swab sample, and clipped into a microcentrifuge tube with Zymo DNA/RNA Shield. Field blanks were subjected to all downstream sample processing and analysis steps.

#### Field lab processing

All household samples were transported back to the local field laboratory in coolers on ice, and sample processing was conducted the same day within approximately 4 hours of sample collection. Swab samples did not require additional processing because swabs were placed directly in Zymo DNA/RNA Shield in the field. Domestic water, drinking water, and child and adult hand rinse samples were filtered through 47 mm 0.45 μm-pore-size polycarbonate filters (Sigma Millipore) using an electric pump. For soil samples, 30 g of unsieved soil was added to a sterile polypropylene bottle with 270 mL PBS and shaken vigorously for two minutes. Soil was allowed to settle for 30 s, then the PBS was decanted into a second polypropylene bottle. The supernatant was filtered using 47-mm 0.45 μm-pore-size polycarbonate filters. The membrane filtration unit was flame-sterilized between each filtered sample, and flame-sterilized forceps were used to handle filters. All filters were stored in 2 mL microcentrifuge tubes containing 1 mL Zymo DNA/RNA Shield. For food samples, 90 mL of PBS was added to 10 g of food in a Whirl-Pak bag and mixed by hand for 2 minutes. 1 mL of diluted and homogenized food (equivalent to 0.1 g of the original food sample) was added to a sterile 2 mL microcentrifuge tube containing 1 mL Zymo DNA/RNA Shield. All samples were stored in at ambient temperature in the field laboratory and during transport to USFQ. Samples were frozen at −80 °C at USFQ prior to DNA extractions.

#### MST qPCR assays

Eight MST qPCR assays were selected for performance validation with respect to identifying fecal contamination from the following fecal sources: humans (HF183, HumM2)^48,49^, birds (AV4143, GFD)^49,50^, dogs (DG37)^51^, pigs (Pig2Bac)^52^, and ruminants (Rum2Bac)^53^. We also tested the performance of a general Bacteroidales fecal marker (GenBac3)^54^ across human and animal fecal sources. HF183, HumM2, AV4143, DG37, and GenBac3 assays were run using an ABI 7500 Real-Time PCR System (Thermo Fisher Scientific; Waltham, US). GFD, Rum2Bac, and Pig2Bac assays were run using a Bio-Rad CFX96 Touch Real-Time PCR Detection System (Bio-Rad Laboratories; California, US). TaqMan performance validation assays were conducted in 25 µL reactions containing 12.5 µL of 1X TaqMan™ Environmental Master Mix (version 2.0; Thermo Fisher Scientific) and 5 µL of DNA template. The GFD SYBR performance validation assay contained 12.5 µL of Power SYBR-Green Master Mix (Thermo Fisher Scientific) and 5 µL of DNA template per reaction. Primer and probe concentrations and cycling conditions are reported in Table 1 for each assay.

**TABLE 1:**
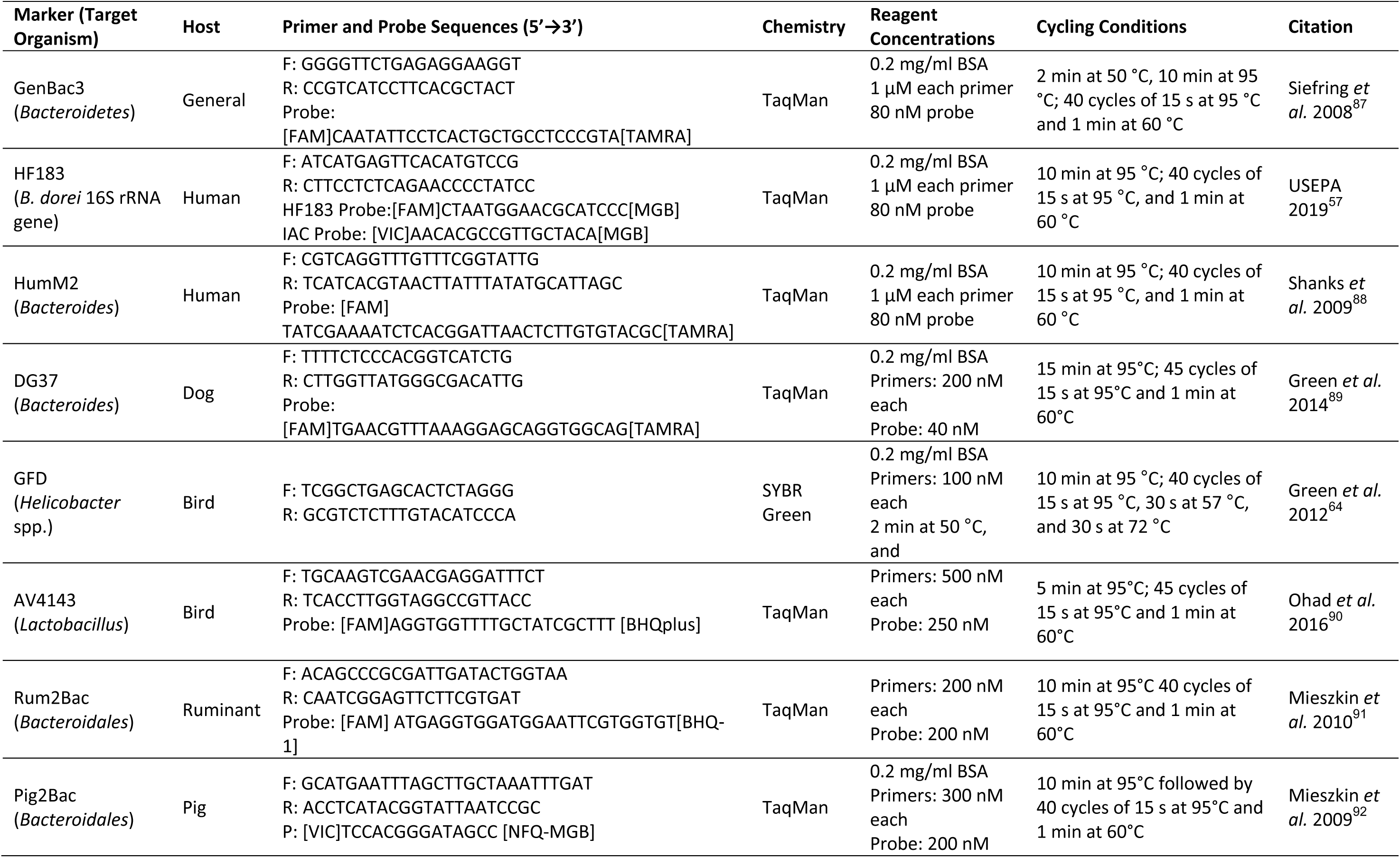
Quantitative PCR assay primers, probes, and reaction conditions.

Marker detection and concentration in animal feces, human feces, and human sewage samples was determined by interpolation to a standard curve, and both standard curve and unknown qPCR reactions were run in triplicate. Standard curves were run alongside unknowns on all reaction plates and were generated using gBlock gene fragments (Integrated DNA Technologies, Coralville, IA) containing the target sequence at ten-fold dilutions ranging from 5 to 10^6^ gene copies per reaction for all markers except GenBac3, where we used dilutions ranging from 10 to 10^7^ gene copies per reaction. Standard curves for the validation study performed within the recommended MIQE guidelines^55,56^ with efficiencies between 90 % and 110 % and R^2^ values > 0.97 (Table S4). Limits of detection (LOD) were defined as the lowest amount of template where all triplicate reactions of the standard were detected for each assay run and were either 5 or 10 gene copies for all assays, depending on the run. Limits of quantification (LOQ) were defined as the lowest concentration that was accurately quantified with an acceptable level of uncertainty and was quantified as: CT_LOQ_=CT_LOD_ – 2(σ_LOD)_. Results were quantified if all three triplicate reactions amplified, fell within 2 Ct values of each other, and were above the level of the lowest standard where all triplicate reactions were detected. If zero, one, or two wells amplified for triplicate reactions, the result was determined to be below the limit of detection (BDL). If triplicate reactions were positive but the amplification occurred after the Ct value for the least concentrated standard, the result was considered detectable but not quantifiable (DNQ). Inhibition of qPCR amplification was assessed with an inhibition amplification control in the HF183 reaction for each sample according to guidelines in USEPA method 1696^57^, and no inhibition of qPCR amplification was detected during performance validation assays. Negative controls including field blanks, lab blanks, extraction blanks and non-template controls for each instrument run were negative or below the detection limit for each assay.

MST assays were validated by assessing their presence in target and non-target animal and/or human samples collected in the study region of northwestern Ecuador, as well as the concentration of each marker in target and non-target fecal samples. Sensitivity was calculated for all candidate assays as the frequency of *true positives* (marker detected in feces from the intended host) divided by the sum of true positives plus *false negatives* (marker not detected in feces from the intended host). Specificity was calculated as the frequency of *true negatives* (marker detected in feces from non-target hosts) divided by the sum of true negatives and *false positives* (marker detected in feces from non-target hosts). A threshold of >80% sensitivity and specificity was applied for determining adequate marker performance, as has been described in previous studies^58,59^. Candidate MST assays with the best performance for each target were used to assess sources of fecal contamination in household samples.

#### Analysis of household samples

##### DNA extractions

DNA was extracted from all household samples using the ZymoBIOMICS DNA Miniprep kit. Household samples and the Zymo DNA/RNA Shield they were stored in were transferred into the provided bead lysis tubes and bead beat at maximum speed on a Bio FastPrep-24 instrument (MP Biomedical) for 3 minutes. Following bead beating, 300 μL supernatant was moved to a new tube with 600 μL DNA/RNA lysis buffer and processed according to the manufacturer’s protocol. Purified DNA for all extractions was eluted with 50 μL elution buffer and immediately stored at −80 °C. Extracted DNA was hand carried from USFQ to USF on gel ice packs as described above for samples from MST validation study, where they were stored at −80°C. DNA concentrations were measured both before freezing at USFQ and prior to qPCR analyses at USF using a Qubit fluorometer.

##### Detection of MST markers in household samples

MST markers in DNA extracted from household samples were detected by qPCR at USF. The following quantitative PCR (qPCR) assays were selected to measure MST markers in DNA extracts from household samples: GenBac3 (general fecal marker)^54^, HF183 (human), DG37 (dog), GFD (avian), Rum2Bac (ruminant), and Pig2Bac (pig). These markers were chosen based on their performance (sensitivity and specificity) during the MST validation study.

All qPCR assays for household samples were run and evaluated using the same laboratory equipment, reagents, reaction conditions, controls, and marker detection and concentration analysis methods as described for the MST validation study (Table 1), except that unknown household samples were run in duplicate and standards were run in triplicate. Assay performance for household sample qPCR runs, including mean efficiencies, slopes, y-intercepts, and R^2^ values are reported in Table S5. Household sample qPCR data were calculated as gene copies per 100 g (food and soil), gene copies per pair of hands (child and adult hand rinses), gene copies per 100 cm^2^ surface area (floors, dish drying and food preparation surfaces), and gene copies per object (keys, toys, cell phones, TV remotes).

### Statistical analyses

Statistical analyses were performed in R version 4.4.2, and visualizations were generated with the R package ggplot2^60^. Statistical significance for all analyses was defined as *p*<0.05. R code is available at https://github.com/kjojess/ECoMiD-household-MST-markers. Samples that were DNQ were considered positive for prevalence analyses and samples that were BDL were considered negative.

#### MST marker prevalence

Differences in marker prevalence were analyzed using Fisher’s exact tests, which are appropriate for count data with small frequencies. For each marker, we ran a global test across all sample types using the command “fisher_test” from the R package rstatix^61^. Because the contingency tables were larger than 2×2, we set the “simulate.p.value” argument set to ‘TRU’ and used 1e4 simulations. For markers with significant *p*-values in the global tests, we used the “fisher.multcomp” function from the R package RVAideMemoire^62^ to conduct pairwise post hoc comparisons between sample types with a Benjamani-Hochberg (BH) correction for multiple testing.

For floor samples and child and adult handwashes, the sample types with the highest proportion of samples that were positive for host-associated markers, we also tested for associations between MST detections and sample characteristics, again using Fisher’s exact tests with 1e4 simulations. For floor samples, we tested for differences in marker detections for various floor types (cement, carpet, vinyl/asphalt, ceramic tiles, wooden boards, or other) and whether the household reported floor cleaning within the last 24 hours (yes or no). For child and adult handwashes, we tested for differences in marker detections by observed hand and nail cleanliness (clean or dirty nails and hands) and reported time since last hand wash (less than 1 hour, 1 hour, less than 2 hours, or more than 2 hours).

Given the role of caregivers in maintaining the household environment and caring for young children, we hypothesized that detection of the human-associated marker HF183 on adult hands would be associated with increased risk of detection in other household sample types. We used Poisson regression with a log link and generalized estimating equations (GEE) to estimate to estimate risk ratios (RRs) and corresponding 95% confidence intervals (CIs) for the association between HF183 detection on adult hands (predictor variable) and HF183 detection in other sample types from the same household (outcome variables). We used an exchangeable correlation structure and clustered by household ID to account for intra-household correlation in the GEE models. Each sample type was analyzed in a separate model, restricted to households in which both the adult hand wash and comparison sample type were collected. We excluded comparisons in which HF183 detection on adult hands perfectly predicted detection in the comparison sample type, resulting in very large risk ratios and CIs (e.g., doorknobs, toys, domestic water, and cellphones). Models were run using the R package geepack^63^.

#### MST marker concentration

We plotted marker concentrations of general, human, and animal-associated MST targets to visually compare differences in marker concentration within each sample type. However, we did not test for between-sample differences in host-associated marker concentration because prevalence of detection for human and animal targets was low (<50% of total samples). Only samples with concentrations >LOQ were plotted. We did test for differences in marker concentration across sample types for the GenBac3 general fecal marker, the only MST target with detections in >50% of samples. For these statistical analyses, samples where GenBac3 was DNQ were assigned the value of the limit of detection (10 copies) and samples where GenBac3 was BDL were assigned a value of half the limit of detection (5 copies). GenBac3 results for soil and food samples were not included because GenBac3 was infrequently detected (<50% prevalence) in these sample types. Shapiro Wilks testing indicated data were non-normal, and a nonparametric Kruskal-Wallis test was used to determine whether median GenBac3 marker concentrations varied by sample type. Pairwise post-hoc Dunn’s tests with a BH correction for multiple testing were performed to identify which sample types were different from one another.

## RESULTS

### MST validation study

Results for the eight microbial source tracking markers we evaluated for use in the study region of northwestern Ecuador are presented in Table 2. Sensitivity/specificity and target/non-target mean abundance data are also presented graphically in Figure S1 and marker percent detection and concentration across target and non-target samples are shown in Figure S2. Samples that were DNQ were considered present and samples that were BDL were considered absent for sensitivity and specificity calculations and prevalence analyses. Samples that were DNQ or BDL were not included in concentration calculations.

**TABLE 2:** MST marker validation study summary.

| Marker | Host | Total samples tested<sup>1</sup> | Target samples tested | Non-target samples tested | Sensitivity | Specificity | Mean concentration in target samples<sup>2</sup> (stdev) | Mean concentration in non-target<sup>1</sup> samples (stdev) |
| --- | --- | --- | --- | --- | --- | --- | --- | --- |
| GenBac | General | 112 | 112 | na | na | na | 10.19 (1.82) | na |
| HF183 | Human | 112 | 7 feces<br>5 sewage | 100 | 100% | 86% | 10.81 (1.17) (feces)<br>8.09 (1.37) (sewage) | 7.39 (2.63) |
| HumM2 | Human | 112 | 7 feces<br>5 sewage | 100 | 92% | 86% | 8.77 (0.71) (feces)<br>6.45 (0.85) (sewage) | 6.6 (1.64) |
| DG37 | Dog | 112 | 21 | 91 | 76% | 92% | 7.17 (1.48) | 6.09 (0.86) |
| AV4143 | Bird | 112 | 28 | 84 | 96% | 80% | 5.17 (1.48) | 4.09 (0.86) |
| GFD | Bird | 112 | 28 | 84 | 82% | 96% | 7.18 (1.50) | 6.80 (0.96) |
| Pig2Bac | Pig | 112 | 17 | 95 | 100% | 67% | 10.44 (1.48) | 5.65 (1.07) |
| Rum2Bac | Ruminant | 112 | 14 | 98 | 100% | 88% | 11.47 (0.22) | 5.34 (1.06) |

<sup>1</sup>Encompasses fecal samples from $n=6$ cats, $n=15$ chickens, $n=14$ cows, $n=21$ dogs, $n=11$ ducks, $n=14$ horses, $n=2$ parrots, $n=17$ pigs, and $n=7$ humans, as well as $n=5$ composite human sewage samples; parrot samples were considered target samples for both AV4143 and GFD MST assays.
<sup>2</sup>log gene copies per 100 g feces or 100 mL sewage.

We found that GenBac3, a marker for general fecal contamination, was present at high concentrations in all human and animal samples tested (10.19 mean log_10_ gene copies per 100 g feces or 100 mL sewage). Two human markers were tested, HF183 and HumM2. HF183 had 100% sensitivity and HumM2 had 92% sensitivity for human samples. Both markers were detected at relatively high concentrations in human samples, with 10.81 and 8.09 mean log_10_ gene copies per 100 g feces or 100 mL sewage for HF183 and HumM2, respectively. Both had 86% specificity; when either marker was detected in a non-target sample it was at approximately three orders of magnitude lower mean concentrations than in human feces or sewage. We selected HF183 as the better-performing marker for human-associated fecal contamination in the study region because it had better sensitivity and was detected at higher concentrations in human feces and sewage than HumM2.

Of the mammalian markers tested, Rum2Bac demonstrated the best overall performance, with 100% sensitivity and 88% specificity. It was present at high concentrations in cow feces (10.44 mean log₁₀ copies per 100 g feces) and at substantially lower concentrations in non-target samples (5.65 mean log₁₀ copies per 100 g feces or 100 mL sewage). Notably, Rum2Bac was not detected in any human feces or sewage samples tested. Pig2Bac was detected in all pig fecal samples tested (100% sensitivity), but exhibited low specificity (67%), amplifying in more than half of cow, horse, duck, and cat samples. Despite this cross-reactivity, Pig2Bac concentrations were markedly higher in pig feces (11.47 mean log₁₀ copies per 100 g feces) compared to non-target samples (5.65 mean log₁₀ copies). The dog-associated marker DG37 showed 76% sensitivity and 91% specificity for dog feces. In addition to its presence in 76% of dog fecal samples, DG37 was detected in sewage, chickens, cats, and one cow sample. However, the concentration of DG37 was approximately tenfold higher in dog feces (mean: 7.17 log₁₀ copies per 100 g feces) than in non-target human and animal samples (mean: 6.09 log₁₀ copies). Although Pig2Bac and DG37 did not meet our predefined threshold of >80% for both sensitivity and specificity, we included them in household sample analysis due to their substantially higher concentrations in target versus non-target feces.

We tested two avian markers, GFD and AV4143, both of which are frequently used as general bird markers^49,64^. AV4143 had 96% specificity for target duck, chicken, and parrot samples, but only 80% specificity. It was frequently detected at high concentrations in non-target feces, especially those from humans, cats, and dogs. GFD on the other hand had higher specificity (96%) but lower sensitivity (82%). GFD was detected in 100% of chicken samples and 72.73% of duck samples but was not detected in parrot samples. Chickens and their feces are common in and around households in the study area, so we selected GFD as the more relevant and better-performing bird marker for this study because of its specificity for domestic poultry and high prevalence in chicken feces.

### MST markers in household samples

We selected GenBac3, HF183, DG37, GFD, Pig2Bac, and Rum2Bac to evaluate household fecal contamination from general, human, dog, bird, pig, and ruminant sources, respectively. In total, we evaluated *n=*585 samples from *n*=58 households. MST marker prevalence and abundance in various sample matrices collected from household are summarized in Figures 2 and 3.

**Figure 2:**
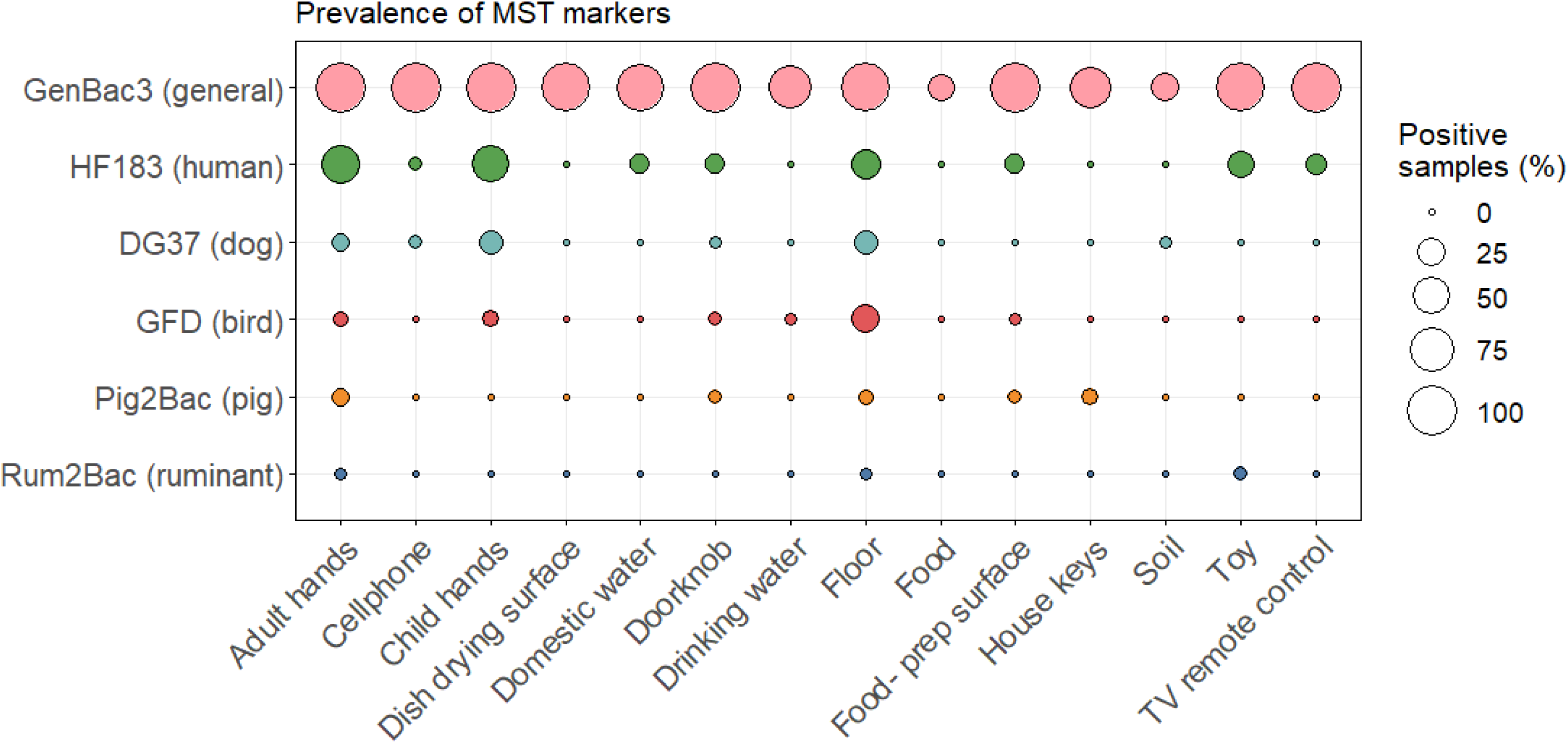
Prevalence of MST markers by household sample type.

**Figure 3:**
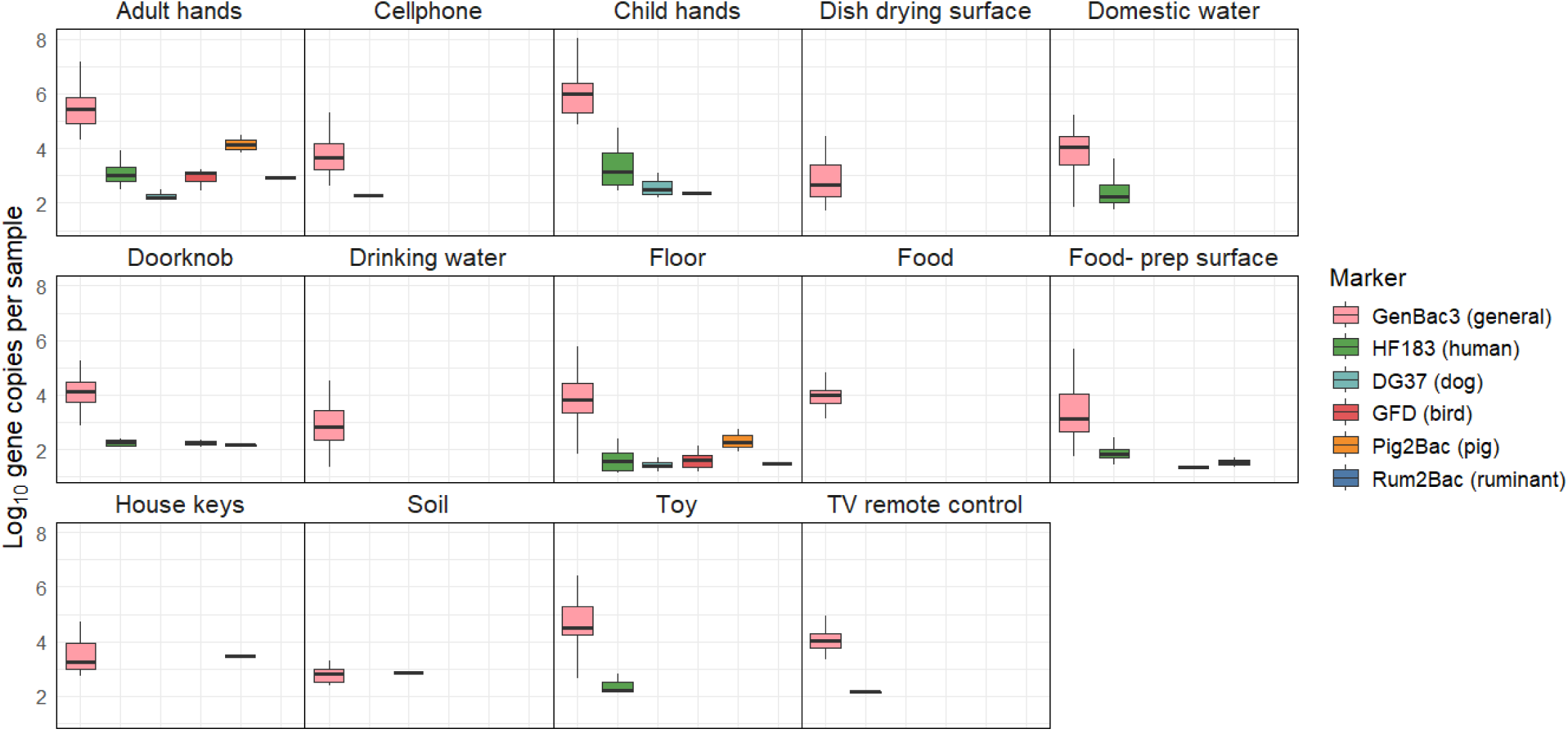
Concentration of MST markers by sample type. Only samples with Ct values >LOQ are included. Household sample qPCR data were calculated as log_10_ gene copies per 100 g (food and soil), gene copies per pair of hands (child and adult hand rinses), gene copies per 100 cm^2^ surface area (floors, dish drying and food preparation surfaces), and gene copies per object (keys, toys, cell phones, TV remotes).

GenBac3, the marker for general fecal contamination, was detected in 78% (456/585) of household samples. It had high prevalence (>50%) in all sample types, except for food and soil. GenBac3 prevalence was different between sample types (Fisher’s exact test *p*<0.001, see Table S6 for *p*-values of between-sample type comparisons). Differences in GenBac3 concentrations were also significantly different between sample types (Kruskal-Walis *p*<0.001); see supplementary Table S7 for a summary of between-sample type results of posthoc Dunn’s testing. We detected the highest concentrations on child and adult and adult hands (6.75 and 6.18 mean log_10_ copies per pair of hands for child and adult hands, respectively) and lowest concentrations in drinking water (3.68 mean log_10_ copies per 100 mL) and on dish drying surfaces (3.56 mean log_10_ copies per 100 cm^2^) (Figure S3).

Human fecal contamination, indicated by detection of HF183, was present in 15% (90/585) of household samples. HF183 was most frequently detected in adult and child hand rinse samples (53% and 48%, respectively), on floors (29%), and on child toys (23%). Animal-associated markers were less frequently detected. GFD, associated with fecal contamination from poultry, found in 4.1% (24/585) of household samples, was the most frequently detected animal marker. We detected GFD in one quarter (25%) of floor samples. DG37, associated with fecal contamination from dogs was detected in 3.6% of household samples. Pig2Bac and Rum2Bac were the least frequently detected animal-associated markers, and indicated fecal contamination from pigs and ruminants, respectively, in 2.0% and 0.52% of household samples. Overall, animal markers were most common on adult hands, child hands, and floors, which were 17%, 22%, and 34% positive for at least one animal marker, respectively. There were no detections of human or animal-associated markers on dish drying surfaces or in food samples. Host-associated markers were infrequently detected in drinking water (one detection of GFD) and soil (one detection of DG37). Statistical testing revealed that prevalence of HF183, DG37, and GFD was significantly different between sample types (*p*<0.001, see Tables S8-S10 for *p*-values of between-sample type marker prevalence comparisons), but there were no differences between sample types in detections of Rum2Bac or Pig2Bac. Despite low prevalence, animal markers were found at relatively high concentrations, comparable to those observed for the human HF183 marker, when detected.

There was no significant relationship between sample characteristics for floor samples (floor material, whether floors were cleaned in the previous 24h) or child/adult hand rinse samples (observed hand and nail cleanliness, time since last handwash) and detection of any MST marker. Detection of the human-specific marker HF183 on adult hands was strongly associated with HF183 detection in other household sample types. In households where adult hands tested positive for HF183, the risk of detection on child hands was over four times higher (RR: 4.27, 95% CI: 1.49–12.20), and the risk of detection in floor samples was nearly seven times higher (RR: 6.77, 95% CI: 1.70–27.04), compared to households where adult hands were negative. Elevated risk of HF183 detection was also observed for food-preparation surfaces, though wide confidence intervals rendered this association statistically non-significant (RR: 4.50, 95% CI: 0.56-36.13). For doorknobs, toys, domestic water, and cellphones, HF183 was only detected when adult hands were also positive, indicating perfect co-occurrence. Risk ratios for these sample types are not reported due to model instability. These results are shown in Figure 4, with model results summarized in Table S11.

**Figure 4.**
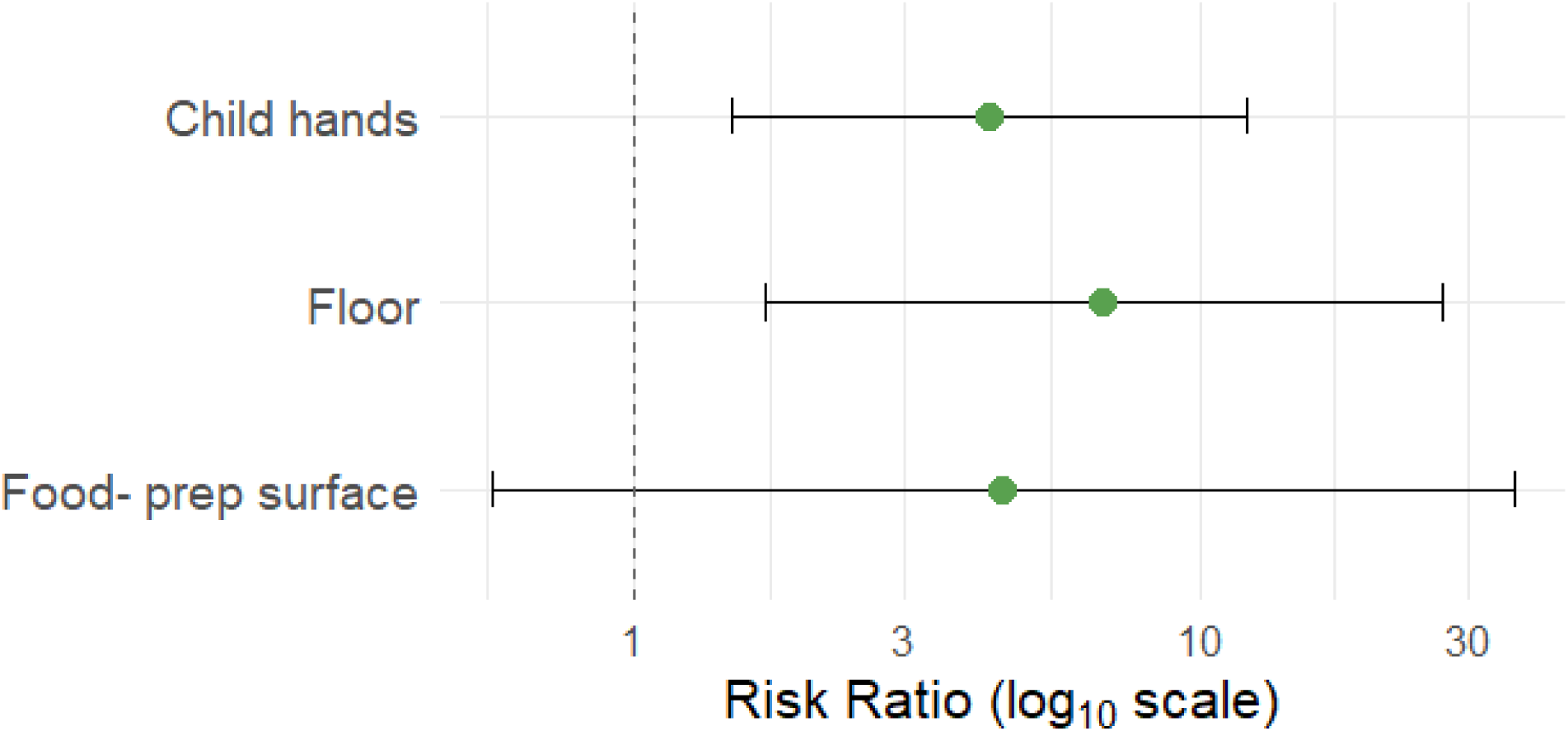
Risk ratios and 95% confidence intervals (CIs) for the association between detection of the human-associated marker HF183 on adult hands and HF183 detection in other sample types within the same household. Risk ratios were calculated using Poisson regression models with robust standard errors. Risk ratios for sample types where HF183 detection was perfectly concordant with adult hands (e.g., toys, doorknobs, domestic water, and cellphones) are not shown.

## DISCUSSION

In this study, we detected widespread fecal contamination across sample matrices in households that owned animals in Northern Ecuador, with the highest prevalence on hands and floors. General fecal contamination was nearly ubiquitous, while human-associated fecal contamination was detected in 15.4% of all samples and animal-associated fecal contamination was detected in 8.4% of all samples. Child and adult hands and floor samples had the highest frequency of MST marker detections, and detections of the human-associated marker HF183 on adult (caregiver) hands were associated with increased likelihood of finding human fecal contamination in other sample types.

### MST validation study

Prior to applying MST to household settings in Northern Ecuador, we evaluated the sensitivity and specificity of MST markers using human and animal feces and human sewage samples. The general fecal marker GenBac3 performed consistently well, with detection at high concentrations across all host fecal and sewage samples. Among human-associated markers, HF183 outperformed HumM2 due to its higher sensitivity and higher concentrations in human feces and sewage. For animal markers, Rum2Bac (ruminants) and GFD (birds) had strong specificity and relevance to local exposures. Performance of the human, avian, and ruminant markers was comparable to or exceeding that reported in previous MST validation studies conducted in LMICs^21,23,24,43,65–69^. In contrast, the dog-associated marker DG37 and the pig-associated marker Pig2Bac showed lower specificity than previously reported. For example, while DG37 had 85% sensitivity and 100% specificity in the U.S.-based laboratory that developed the assay^51^, we observed 76% sensitivity and 92% specificity in Northern Ecuador. Pig2Bac had high specificity (≥90%) for pig feces in studies conducted in Peru, India, Nepal, and China^24,43,65,69^, but only 67% specificity in our study. These differences reflect cross-species cross-reactivity for these markers and are attributable to regional variations in gut microbiota composition among animal hosts^70^. Despite these limitations, both DG37 and Pig2Bac were present at much higher concentrations in their respective target samples than in non-target samples, and we included them for source attribution in this study.

### Insights into sources of fecal contamination

General fecal contamination was widespread, with the GenBac3 marker detected at high prevalence across all household sample types. The human-associated HF183 marker was the most frequently detected host-specific marker, present in 15.4% of samples, and most commonly found on adult and child hands, followed by floors and toys. In contrast, animal-associated MST markers were detected in only 8.4% of samples, surprisingly low prevalence given that animal ownership was a criterion for inclusion and prior observations describing frequent presence of animals and their feces in and around households in the study communities^71^. However, when detected, animal markers were present at high concentrations, suggesting that even infrequent contamination events may present significant health risks.

Despite infrequent detection of animal markers when considering all sample types, we observed relatively high prevalence of animal markers on floors (33.9%) and child and caregiver hand rinses (22.6 and 17.0%, respectively). We recently reported that there are myriad opportunities for children to be exposed to animals and their feces in household environments regardless of whether the household owned animals^47^. For example, in some study communities, it is common practice to allow animals to roam freely to forage for food, which is perceived as beneficial to animal health and reduces the cost of feed. This behavior contributes to the pervasive presence of animal feces on floors and in other household spaces, increasing the potential for child exposure. Studies in high-income country settings

Among the animal markers, the dog-associated DG37 and bird-associated GFD were detected more frequently than markers for pigs or ruminants. This distribution aligns with household animal ownership patterns: dogs and poultry were the most commonly owned animals among study households, followed by cats (who tend to bury their feces and for which no validated MST marker exists), pigs, and cattle. Prior research in this region indicates that pigs and cattle are typically penned and kept away from household living areas, whereas dogs and chickens roam freely^71^. We also frequently observed dog and chicken feces in and around household environments during data collection for this and previous studies^71,72^.

Frequent MST markers on child and adult caregiver hands and on household floors indicates that these may be important reservoirs of fecal contamination. Previous findings in LMICs have also highlighted hands and household surface as vectors for household-level transmission of fecal contamination and enteric pathogens. For example, studies in Bangladesh and Mozambique reported frequent detection of HF183 and other human-associated MST markers on hands and household surfaces^21,23^. Research studies in Bangladesh and Honduras showed that reduced FIB detection and concentration on toys were associated with improved WaSH conditions^7,73^, though a study in Kenya found no impact of a WaSH intervention on FIB contamination of child toys^74^. Fecal contamination has also been previously measured on floors in LMIC settings, with studies in Peru showing less FIB contamination on improved flooring types and in households with improved sanitation^75^ and more frequent detections of human, avian, and dog markers on floors than other household surfaces^24^. Contamination of household surfaces, particularly toys and floors, and child hands is concerning given that enteric pathogen infections have the greatest negative health impacts on very young children. We have observed infants playing on household floors and demonstrating developmentally-appropriate infant mouthing behaviors of hands and objects in study communities^72,76^.

Detection of the human-associated HF183 marker on adult hands was significantly associated with its detection on child hands and household floors, suggesting shared contamination pathways between caregivers, children, and domestic surfaces. HF183 was also detected on doorknobs, cellphones, child toys, and in domestic water, but we were unable to statistically evaluate these associations because adult hands tested positive in all corresponding cases. These findings reinforce the idea that caregivers may serve as central nodes in the microbial transfer network within the household environment, facilitating the spread of fecal contamination across people and surfaces. Evidence from other settings supports this hypothesis. In Tanzania, the concentration of fecal indicator bacteria (FIB) on caregiver hands was the strongest predictor of FIB levels in drinking water, with behaviors such as food preparation, recent handwashing, and movement in and out of the home associated with increased contamination. Similarly, a study in Kenya used 16S rRNA gene amplicon sequencing to distinguish microbial communities derived from child versus adult feces and found that caregiver hands were frequently contaminated with child feces. While these findings suggest a key role for caregivers in intra-household contamination dynamics, the cross-sectional design of our study limits our ability to determine directionality. It is possible that adult hands became contaminated through contact with already contaminated surfaces, such as floors or shared household objects. Nonetheless, the frequent detection of human fecal contamination on caregiver hands highlights a direct and potentially high-risk exposure pathway for young children.

Though they have not employed MST methods, several studies conducted in high income countries (HICs) have reported the presence of FIB and pathogens on hands and surfaces in households and other indoor settings^77^. For example, a study examining kitchen surfaces in Philadelphia, PA found fecal coliforms in 44% and enteric pathogens in 45% of households^78^. These results suggest that fecal contamination in household settings is not unique to LMICs, though the contamination pathways, microbial sources, and health risks may differ substantially between LMIC and high-income settings. A recent systematic review and meta-analysis found that rates of *E. coli* detection in hand rinse samples were significantly higher in LMICs compared to HICs^79^, consistent with the disproportionate burden of enteric disease in LMICs^80^.

We found a high prevalence of HF183 in households, suggesting widespread human fecal contamination and highlighting the potential importance of addressing human-associated sources to reduce pathogen exposure. We did not attempt to link MST markers to household water, sanitation, or hygiene conditions. However, recent studies have found that sanitation interventions had no impact on the prevalence of human- or animal-associated MST markers in household or environmental samples^40,81^. One exception was a sanitation intervention study that focused on improving management of animal and child feces, where there was reduced incidence of the ruminant-associated marker BacR in water samples and general fecal contamination in soil in Bangladesh^21^. These null effects are consistent with the small or null health effects reported in recent WaSH trials that suggest that basic WaSH interventions do not contain sources of human fecal contamination or adequately reduce exposure to environmental enteropathogens^81^. Continued research is needed to understand which household environmental conditions and interventions aimed at improving them can meaningfully reduce environmental fecal contamination and associated child health outcomes.

### Limitations

We observed relatively low prevalence of host-associated MST markers in food and drinking water, matrices that are widely recognized as important reservoirs for enteric pathogen transmission. However, the general fecal marker GenBac3 was detected in 70.1% of drinking water samples and 21.1% of food samples, indicating frequent contamination with fecal material from undetermined sources. The limited detection of host-specific markers in food may be partly attributable to methodological constraints, as DNA extractions were conducted on a very small subsample (∼0.1 g) of the original food material, potentially limiting sensitivity. Similarly, we detected MST markers infrequently in soil samples, in contrast to other studies that have reported frequent detection of *E. coli* and human- and animal-associated MST markers in soil in LMIC settings^12,44,82,83^. Our soil processing protocol was adapted from a method originally developed for MST marker recovery in beach sand^84^, which may not have worked well for the more the heterogeneous and compacted soils sampled in this study.

Differences in sample collection, processing, and extraction methods across matrices make direct comparisons between sample types challenging. As such, cross-sample comparisons, particularly for concentration data, should be interpreted with caution. Despite these limitations, the inclusion of multiple environmental sample types and MST markers offers a more complete view of potential fecal exposure pathways within the household environment.

The MST qPCR assays used in this study detect host-associated bacterial DNA markers and do not provide information on viability, an important consideration when assessing whether detection is indicative of the presence of live, infectious pathogens^85^. Additionally, all markers targeted bacteria. However, bacterial, viral, and parasitic enteric pathogens differ in their environmental persistence^86^. Finally, although our analysis focused on household-level contamination, community-level exposures to animals and their feces through shared outdoor spaces and environmental fecal contamination from roaming animals may also contribute to enteric pathogen transmission and warrant further investigation^72^.

### Conclusions

This study presents one of the most comprehensive assessments to date of LMIC household fecal contamination sources, incorporating multiple MST markers across diverse environmental sample types. Compared to many previous MST studies in LMIC settings, our inclusion of six MST markers across ten diverse environmental sample types, including hands, household surfaces, soil, water, and prepared food, provides a more comprehensive assessment of fecal contamination within domestic environments. This breadth allows for deeper insights into potential exposure pathways and microbial transmission dynamics, particularly in complex household settings. In addition, identifying the sample types where we are most likely to detect general and host-associated fecal contamination (floor and hand rinse samples) will help guide sampling strategies for future studies.

Our findings highlight child and adult caregiver hands and household floors as reservoirs of both human and animal fecal contamination, underscoring their potential role in enteric pathogen transmission. The association between HF183 detection on adult hands and contamination on child hands and floors suggests that caregiver hygiene may influence intra-household contamination patterns. While animal-associated markers were less prevalent overall, their detection at high concentrations, particularly for dogs and chickens, indicates that zoonotic exposures may contribute meaningfully to environmental contamination and enteric disease risk. These results underscore the importance of integrated interventions that address both human and animal fecal sources and highlight the potential utility of MST methods for identifying priority exposure pathways. Future research should examine how contamination patterns shift over time and assess links between household contamination, comprehensive measurements of animal exposures and WaSH conditions, and child health outcomes to inform targeted strategies for interrupting fecal-oral transmission of enteric pathogens.

## Acknowledgements

We are grateful to all the participants who enrolled in this study and provided us access to their homes. This research would not have been possible without our dedicated local field staff who helped us identify eligible households and assisted in sample collection. Thank you to Drs. Caitlin Hemlock and Berry Brosi for advice on the statistical analyses. This work was funded by the National Institutes of Health (R01AI137679 and R01AI162867) and supported under award numbers 5T32ES007032-37 (KJJ), 5T32ES012870 (KJJ), and K01AI145080 (GOL). The content is solely the responsibility of the authors and does not necessarily represent the official views of the National Institutes of Health.

## Citations

1. Wagner, E; Lanoix J. Excreta Disposal for Rural Areas and Small Communities. vol. 39 (Monograph Series, World Health Organization).

2. Mara, D., Lane, J., Scott, B. & Trouba, D. Sanitation and Health. PLOS Medicine 7, e1000363 (2010).

3. Guerrant, R. L., Deboer, M. D., Moore, S. R., Scharf, R. J. & Lima, A. A. M. The Impoverished Gut - A Triple Burden of Diarrhoea, Stunting and Chronic Disease. Nature Reviews Gastroenterology and Hepatology vol. 10 (Nature Publishing Group, 2013).

4. Investigators, M.-E. N. Early childhood cognitive development is affected by interactions among illness, diet, enteropathogens and the home environment: findings from the MAL-ED birth cohort study. BMJ Global Health 3, e000752. PMCID: PMC6058175. (2018).

5. Pickard, J. M., Zeng, M. Y., Caruso, R. & Núñez, G. Gut microbiota: Role in pathogen colonization, immune responses, and inflammatory disease. Immunological Reviews 279, 70–89 (2017).

6. Clasen, T., et al. Effectiveness of a rural sanitation programme on diarrhoea, soil-transmitted helminth infection, and child malnutrition in Odisha, India: a cluster-randomised trial. The Lancet Global Health 2, e645–e653 (2014).

7. Humphrey, J. H. et al. Independent and combined effects of improved water, sanitation, and hygiene, and improved complementary feeding, on child stunting and anaemia in rural Zimbabwe: a cluster-randomised trial. The Lancet Global Health 7, e132–e147 (2019).

8. Patil, S. R. et al. The Effect of India’s Total Sanitation Campaign on Defecation Behaviors and Child Health in Rural Madhya Pradesh: A Cluster Randomized Controlled Trial. PLOS Medicine 11, e1001709 (2014).

9. Sclar, G. D. et al. Assessing the impact of sanitation on indicators of fecal exposure along principal transmission pathways: A systematic review. International Journal of Hygiene and Environmental Health 219, 709–723 (2016).

10. Knee, J., et al. Effects of an urban sanitation intervention on childhood enteric infection and diarrhea in Maputo, Mozambique: A controlled before-and-after trial. eLife 10, e62278 (2021).

11. Fuhrmeister, E. R. et al. Effect of Sanitation Improvements on Pathogens and Microbial Source Tracking Markers in the Rural Bangladeshi Household Environment. Environ. Sci. Technol. 54, 4316–4326 (2020).

12. Ercumen, A. et al. Animal Feces Contribute to Domestic Fecal Contamination: Evidence from E. coli Measured in Water, Hands, Food, Flies, and Soil in Bangladesh. Environ Sci Technol 51, 8725–8734 (2017).

13. Penakalapati, G. et al. Exposure to animal feces and human health: A systematic review and proposed research priorities. Environ. Sci. Technol. 51, 11537–11552 (2017).

14. Delahoy, M. J., et al. Pathogens Transmitted in Animal Feces in Low- and Middle-Income Countries. International Journal of Hygiene and Environmental Health vol. 221 (Elsevier GmbH, 2018).

15. Prendergast, A. J. et al. Putting the “A” into WaSH: a call for integrated management of water, animals, sanitation, and hygiene. The Lancet Planetary Health 3, e336–e337 (2019).

16. Ahmed, W. et al. Marker genes of fecal indicator bacteria and potential pathogens in animal feces in subtropical catchments. Science of The Total Environment 656, 1427–1435 (2019).

17. Kagambèga, A. et al. Prevalence and characterization of Salmonella enterica from the feces of cattle, poultry, swine and hedgehogs in Burkina Faso and their comparison to human Salmonella isolates. BMC Microbiol 13, 253 (2013).

18. Kagambèga, A. et al. Prevalence of diarrheagenic Escherichia coli virulence genes in the feces of slaughtered cattle, chickens, and pigs in Burkina Faso. MicrobiologyOpen 1, 276–284 (2012).

19. Hutchison, M. L., Walters, L. D., Avery, S. M., Munro, F. & Moore, A. Analyses of Livestock Production, Waste Storage, and Pathogen Levels and Prevalences in Farm Manures. Applied and Environmental Microbiology 71, 1231–1236 (2005).

20. Ewers, C., Antão, E.-M., Diehl, I., Philipp, H.-C. & Wieler, L. H. Intestine and Environment of the Chicken as Reservoirs for Extraintestinal Pathogenic Escherichia coli Strains with Zoonotic Potential. Applied and Environmental Microbiology 75, 184–192 (2009).

21. Boehm, A. B. et al. Occurrence of Host-Associated Fecal Markers on Child Hands, Household Soil, and Drinking Water in Rural Bangladeshi Households. ACS Publications https://pubs.acs.org/doi/pdf/10.1021/acs.estlett.6b00382 (2016) doi:10.1021/acs.estlett.6b00382.

22. Carron, M. et al. Campylobacter, a zoonotic pathogen of global importance: Prevalence and risk factors in the fast-evolving chicken meat system of Nairobi, Kenya. PLoS Negl Trop Dis 12, e0006658 (2018).

23. Holcomb, D. A. et al. Human fecal contamination of water, soil, and surfaces in households sharing poor-quality sanitation facilities in Maputo, Mozambique. International Journal of Hygiene and Environmental Health 226, 113496 (2020).

24. Schiaffino, F. et al. Associations among Household Animal Ownership, Infrastructure, and Hygiene Characteristics with Source Attribution of Household Fecal Contamination in Peri-Urban Communities of Iquitos, Peru. Am J Trop Med Hyg 104, 372–381 (2021).

25. Stoeckel, D. M. & Harwood, V. J. Performance, Design, and Analysis in Microbial Source Tracking Studies. Applied and Environmental Microbiology 73, 2405–2415 (2007).

26. Field, K. G. & Samadpour, M. Fecal source tracking, the indicator paradigm, and managing water quality. Water Research 41, 3517–3538 (2007).

27. Harwood, V. J., Staley, C., Badgley, B. D., Borges, K. & Korajkic, A. Microbial source tracking markers for detection of fecal contamination in environmental waters: relationships between pathogens and human health outcomes. FEMS Microbiology Reviews 38, 1–40 (2014).

28. Capone, D. et al. Quantitative Microbial Risk Assessment of Pediatric Infections Attributable to Ingestion of Fecally Contaminated Domestic Soils in Low-Income Urban Maputo, Mozambique. Environ. Sci. Technol. 55, 1941–1952 (2021).

29. Julian, T. R. et al. Fecal Indicator Bacteria Contamination of Fomites and Household Demand for Surface Disinfection Products: A Case Study from Peru. Am J Trop Med Hyg 89, 869–872 (2013).

30. Gizaw, Z., Yalew, A. W., Bitew, B. D., Lee, J. & Bisesi, M. Fecal indicator bacteria along multiple environmental exposure pathways (water, food, and soil) and intestinal parasites among children in the rural northwest Ethiopia. BMC Gastroenterology 22, 84 (2022).

31. Pickering, A. J. et al. Fecal Indicator Bacteria along Multiple Environmental Transmission Pathways (Water, Hands, Food, Soil, Flies) and Subsequent Child Diarrhea in Rural Bangladesh. Environ. Sci. Technol. 52, 7928–7936 (2018).

32. Barrett, L. R. et al. Beyond borders: A systematic review and meta-analysis of human-specific faecal markers across geographical settings. Critical Reviews in Environmental Science and Technology 55, 447–464 (2025).

33. Harris, A. R. et al. Ruminants Contribute Fecal Contamination to the Urban Household Environment in Dhaka, Bangladesh. ACS Publications https://pubs.acs.org/doi/pdf/10.1021/acs.est.5b06282 (2016) doi:10.1021/acs.est.5b06282.

34. Fuhrmeister, E. R. et al. Predictors of Enteric Pathogens in the Domestic Environment from Human and Animal Sources in Rural Bangladesh. Environ. Sci. Technol. 53, 10023–10033 (2019).

35. Schriewer, A. et al. Human and Animal Fecal Contamination of Community Water Sources, Stored Drinking Water and Hands in Rural India Measured with Validated Microbial Source Tracking Assays. The American Journal of Tropical Medicine and Hygiene 93, 509–516 (2015).

36. Odagiri, M. et al. Human fecal and pathogen exposure pathways in rural Indian villages and the effect of increased latrine coverage. Water Research 100, 232–244 (2016).

37. Malla, B. et al. Validation of host-specific Bacteroidales quantitative PCR assays and their application to microbial source tracking of drinking water sources in the Kathmandu Valley, Nepal. Journal of Applied Microbiology 125, 609–619 (2018).

38. Healy-Profitós, J. et al. Neighborhood diversity of potentially pathogenic bacteria in drinking water from the city of Maroua, Cameroon. Journal of Water and Health 14, 559–570 (2016).

39. Mills, M. et al. Household environment and animal fecal contamination are critical modifiers of the gut microbiome and resistome in young children from rural Nicaragua. Microbiome 11, 207 (2023).

40. Holcomb, D. A. et al. Associations between fecal contamination of the household environment and enteric pathogen detection in children living in Maputo, Mozambique. Preprint at 10.1101/2025.03.11.25323794 (2025).

41. Daly, S. W. et al. Enteric Pathogens in Humans, Domesticated Animals, and Drinking Water in a Low-Income Urban Area of Nairobi, Kenya. Environ. Sci. Technol. 58, 21839–21849 (2024).

42. Zhang, Y., Wu, R., Lin, K., Wang, Y. & Lu, J. Performance of host-associated genetic markers for microbial source tracking in China. Water Research 175, 115670 (2020).

43. Odagiri, M. et al. Validation of Bacteroidales quantitative PCR assays targeting human and animal fecal contamination in the public and domestic domains in India. Science of The Total Environment 502, 462–470 (2015).

44. Boehm, A. B. et al. Occurrence of Host-Associated Fecal Markers on Child Hands, Household Soil, and Drinking Water in Rural Bangladeshi Households. Environ. Sci. Technol. Lett. 3, 393–398 (2016).

45. Jesser, K. J. et al. Environmental Exposures Associated with Enteropathogen Infection in Six-Month-Old Children Enrolled in the ECoMiD Cohort along a Rural–Urban Gradient in Northern Ecuador. Environ. Sci. Technol. 59, 103–118 (2025).

46. Montero, L. et al. Distribution of Escherichia coli Pathotypes along an Urban–Rural Gradient in Ecuador. Am J Trop Med Hyg 109, 559–567 (2023).

47. Ballard, A. M. et al. Multilevel factors drive child exposure to enteric pathogens in animal feces: A qualitative study in northwestern coastal Ecuador. PLOS Glob Public Health 4, e0003604 (2024).

48. Shanks, O. C., Kelty, C. A., Sivaganesan, M., Varma, M. & Haugland, R. A. Quantitative PCR for genetic markers of human fecal pollution. Appl Environ Microbiol 75, 5507–5513 (2009).

49. Ohad, S. et al. The Development of a Novel qPCR Assay-Set for Identifying Fecal Contamination Originating from Domestic Fowls and Waterfowl in Israel. Frontiers in Microbiology 7, (2016).

50. McMinn, B. R. et al. A constructed wetland for treatment of an impacted waterway and the influence of native waterfowl on its perceived effectiveness. Ecological Engineering 128, 48–56 (2019).

51. Green, H. C., White, K. M., Kelty, C. A. & Shanks, O. C. Development of Rapid Canine Fecal Source Identification PCR-Based Assays. Environ. Sci. Technol. 48, 11453–11461 (2014).

52. Mieszkin, S., Furet, J.-P., Corthier, G. & Gourmelon, M. Estimation of Pig Fecal Contamination in a River Catchment by Real-Time PCR Using Two Pig-Specific Bacteroidales 16S rRNA Genetic Markers. Applied and Environmental Microbiology 75, 3045–3054 (2009).

53. Mieszkin, S., Yala, J.-F., Joubrel, R. & Gourmelon, M. Phylogenetic analysis of Bacteroidales 16S rRNA gene sequences from human and animal effluents and assessment of ruminant faecal pollution by real-time PCR. Journal of Applied Microbiology 108, 974–984 (2010).

54. Siefring, S., Varma, M., Atikovic, E., Wymer, L. & Haugland, R. A. Improved real-time PCR assays for the detection of fecal indicator bacteria in surface waters with different instrument and reagent systems. Journal of Water and Health 6, 225–237 (2008).

55. Huggett, J. F. et al. The digital MIQE guidelines: Minimum Information for Publication of Quantitative Digital PCR Experiments. Clin Chem 59, 892–902 (2013).

56. Bustin, S. A. et al. The MIQE Guidelines: Minimum Information for Publication of Quantitative Real-Time PCR Experiments. Clinical Chemistry 55, 611–622 (2009).

57. USEPA. Method 1696: Characterization of Human Fecal Pollution in Water by HF183/BacR287 TaqMan Quantitative Polymerase Chain Reaction (qPCR) Assay. (2019).

58. Barrett, L. R. et al. Beyond borders: A systematic review and meta-analysis of human-specific faecal markers across geographical settings. Crit Rev Environ Sci Technol 55, 447–464.

59. Rytkönen, A. et al. The Use of Ribosomal RNA as a Microbial Source Tracking Target Highlights the Assay Host-Specificity Requirement in Water Quality Assessments. Front. Microbiol. 12, (2021).

60. Pedersen, H. W., Danielle Navarro, and Thomas Lin. Ggplot2: Elegant Graphics for Data Analysis.

61. Alboukadel, K. rstatix: Pipe-Friendly Framework for Basic Statistical Tests. CRAN: Contributed Packages (2019) doi:10.32614/cran.package.rstatix.

62. Hervé: RVAideMemoire: testing and plotting procedures… - Google Scholar. https://scholar.google.com/scholar_lookup?author=M+Herv%C3%A9+&publication_year=2021&title=RVAideMemoire%3A+Testing+and+Plotting+Procedures+for+Biostatistics#d=gs_cit&t=1714685757943&u=%2Fscholar%3Fq%3Dinfo%3AOIU0az5EUp8J%3Ascholar.google.com%2F%26output%3Dcite%26scirp%3D0%26hl%3Den.

63. Højsgaard, S., Halekoh, U. & Yan, J. The R Package geepack for Generalized Estimating Equations. Journal of Statistical Software 15, 1–11 (2006).

64. Green, H. C., Dick, L. K., Gilpin, B., Samadpour, M. & Field, K. G. Genetic Markers for Rapid PCR-Based Identification of Gull, Canada Goose, Duck, and Chicken Fecal Contamination in Water. Appl Environ Microbiol 78, 503–510 (2012).

65. Malla, B. et al. Validation of host-specific Bacteroidales quantitative PCR assays and their application to microbial source tracking of drinking water sources in the Kathmandu Valley, Nepal. Journal of Applied Microbiology 125, 609–619 (2018).

66. Linke, R. B. et al. Assessing the faecal source sensitivity and specificity of ruminant and human genetic microbial source tracking markers in the central Ethiopian highlands. Letters in Applied Microbiology 72, 458–466 (2021).

67. Yahya, M. et al. Comparison of the Performance of Different Microbial Source Tracking Markers among European and North African Regions. Journal of Environmental Quality 46, 760–766 (2017).

68. Symonds, E. M. et al. Microbial source tracking in shellfish harvesting waters in the Gulf of Nicoya, Costa Rica. Water Research 111, 177–184 (2017).

69. Vadde, K. K., McCarthy, A. J., Rong, R. & Sekar, R. Quantification of Microbial Source Tracking and Pathogenic Bacterial Markers in Water and Sediments of Tiaoxi River (Taihu Watershed). Front Microbiol 10, 699 (2019).

70. Senghor, B., Sokhna, C., Ruimy, R. & Lagier, J.-C. Gut microbiota diversity according to dietary habits and geographical provenance. Human Microbiome Journal 7–8, 1–9 (2018).

71. Alban, V., et al. Exploring child exposure to animal feces and zoonotic pathogens in northern coastal Ecuador: A mixed methods study. In Prep.

72. Ballard, A. M. et al. Multilevel factors drive child exposure to enteric pathogens in animal feces: A qualitative study in northwestern coastal Ecuador. PLOS Glob Public Health 4, e0003604 (2024).

73. Vujcic, J. et al. Toys and toilets: cross-sectional study using children’s toys to evaluate environmental faecal contamination in rural Bangladeshi households with different sanitation facilities and practices. Tropical Medicine & International Health 19, 528–536 (2014).

74. Swarthout, J. M. et al. Addressing Fecal Contamination in Rural Kenyan Households: The Roles of Environmental Interventions and Animal Ownership. Environ. Sci. Technol. 58, 9500–9514 (2024).

75. Exum, N. G. et al. Floors and Toilets: Association of Floors and Sanitation Practices with Fecal Contamination in Peruvian Amazon Peri-Urban Households. Environ. Sci. Technol. 50, 7373–7381 (2016).

76. Sosa-Moreno, A. et al. Characterizing Behaviors Associated with Enteric Pathogen Exposure among Infants in Rural Ecuador through Structured Observations. Am J Trop Med Hyg 106, 1747–1756 (2022).

77. Stephens, B. et al. Microbial Exchange via Fomites and Implications for Human Health. Curr Pollution Rep 5, 198–213 (2019).

78. Borrusso, P. A. & Quinlan, J. J. Prevalence of Pathogens and Indicator Organisms in Home Kitchens and Correlation with Unsafe Food Handling Practices and Conditions. Journal of Food Protection 80, 590–597 (2017).

79. Cantrell, M. E. et al. Hands Are Frequently Contaminated with Fecal Bacteria and Enteric Pathogens Globally: A Systematic Review and Meta-analysis. ACS Environ. Au 3, 123–134 (2023).

80. Troeger, C. et al. Estimates of the global, regional, and national morbidity, mortality, and aetiologies of diarrhoea in 195 countries: a systematic analysis for the Global Burden of Disease Study 2016. The Lancet Infectious Diseases 18, 1211–1228 (2018).

81. Mertens, A. et al. Effects of water, sanitation, and hygiene interventions on detection of enteropathogens and host-specific faecal markers in the environment: a systematic review and individual participant data meta-analysis. The Lancet Planetary Health 7, e197–e208 (2023).

82. Fuhrmeister, E. R. et al. Predictors of Enteric Pathogens in the Domestic Environment from Human and Animal Sources in Rural Bangladesh. Environ. Sci. Technol. 53, 10023–10033 (2019).

83. Holcomb, D. A. et al. Human fecal contamination of water, soil, and surfaces in households sharing poor-quality sanitation facilities in Maputo, Mozambique. International Journal of Hygiene and Environmental Health 226, 113496 (2020).

84. Gallard-Gongora, J., Lobos, A., Conrad, J. W., Peraud, J. & Harwood, V. J. An assessment of three methods for extracting bacterial DNA from beach sand. J Appl Microbiol 132, 2990–3000 (2022).

85. Trinh, K. T. L. & Lee, N. Y. Recent Methods for the Viability Assessment of Bacterial Pathogens: Advances, Challenges, and Future Perspectives. Pathogens 11, 1057 (2022).

86. Hopkins, S. R. et al. Environmental Persistence of the World’s Most Burdensome Infectious and Parasitic Diseases. Front Public Health 10, 892366 (2022).

87. Siefring, S., Varma, M., Atikovic, E., Wymer, L. & Haugland, R. A. Improved real-time PCR assays for the detection of fecal indicator bacteria in surface waters with different instrument and reagent systems. Journal of Water and Health 6, 225–237 (2008).

88. Shanks, O. C., Kelty, C. A., Sivaganesan, M., Varma, M. & Haugland, R. A. Quantitative PCR for genetic markers of human fecal pollution. Appl Environ Microbiol 75, 5507–5513 (2009).

89. Green, H. C., White, K. M., Kelty, C. A. & Shanks, O. C. Development of Rapid Canine Fecal Source Identification PCR-Based Assays. Environ. Sci. Technol. 48, 11453–11461 (2014).

90. Ohad, S. et al. The Development of a Novel qPCR Assay-Set for Identifying Fecal Contamination Originating from Domestic Fowls and Waterfowl in Israel. Frontiers in Microbiology 7, (2016).

91. Mieszkin, S., Yala, J.-F., Joubrel, R. & Gourmelon, M. Phylogenetic analysis of Bacteroidales 16S rRNA gene sequences from human and animal effluents and assessment of ruminant faecal pollution by real-time PCR. Journal of Applied Microbiology 108, 974–984 (2010).

92. Mieszkin, S., Furet, J.-P., Corthier, G. & Gourmelon, M. Estimation of Pig Fecal Contamination in a River Catchment by Real-Time PCR Using Two Pig-Specific Bacteroidales 16S rRNA Genetic Markers. Applied and Environmental Microbiology 75, 3045–3054 (2009).

